# Gut microbiota profile in CDKL5 deficiency disorder patients as a potential marker of clinical severity

**DOI:** 10.1101/2023.12.01.569361

**Authors:** Elisa Borghi, Ornella Xynomilakis, Emerenziana Ottaviano, Camilla Ceccarani, Ilaria Viganò, Paola Tognini, Aglaia Vignoli

## Abstract

CDKL5 deficiency disorder (CDD) is a neurodevelopmental condition characterized by global developmental delay, early-onset seizures, intellectual disability, visual and motor impairments. Unlike Rett Syndrome (RTT), CDD lacks a clear regression period. CDD patients frequently encounter gastrointestinal (GI) disturbances and exhibit signs of subclinical immune dysregulation. However, the underlying causes of these conditions remain elusive. Emerging studies indicate a potential connection between neurological disorders and gut microbiota, an area completely unexplored in CDD. We conducted a pioneering study, analyzing fecal microbiota composition in CDD patients and their healthy relatives. Notably, differences in intestinal bacterial diversity and composition were identified in CDD patients. We further investigated microbiota changes based on the severity of GI issues, seizure frequency, sleep disorders, food intake type, impairment in neuro-behavioral features (assessed through the RTT Behaviour Questionnaire – RSBQ), and ambulation capacity.

Our findings hint at a potential connection between CDD, microbiota, and symptom severity. This study marks the first exploration of the gut-microbiota-brain axis in CDD patients. It adds to the growing body of research emphasizing the role of the gut microbiota in neurodevelopmental disorders and opens doors to potential interventions that target intestinal microbes with the aim of improving the lives of CDD patients.

## INTRODUCTION

CDKL5 deficiency disorder (CDD) is a neurodevelopmental disorder caused by mutations in the X-linked cyclin-dependent kinase-like 5 *CDKL5* gene. It leads to severe global developmental delay, early-onset epileptic encephalopathy, intellectual disability, cortical visual impairment, fine and gross motor impairment, hypotonia, sleep disturbances, and hand stereotypies^1^. While some symptoms are similar to Rett Syndrome (RTT), CDD is distinguished by the lack of a regression period and the early seizure onset. Currently, no treatments are available for CDD.

Patients with CDD often experience gastrointestinal (GI) problems^2^, altered growth pattern^3^, and have subclinical immune dysfunction, likely caused by a malfunctioning inflammatory regulatory system^4^. The reason for this dysfunction is not well understood, but it has also been observed in patients with classic RTT. Studies have found that RTT patients display changes in their gut microbiota composition^5,6^, which is the collection of microorganisms that live in the GI tract in equilibrium with their host^7^. This recent research suggests that changes in the gut could be linked to neurological issues. For example, multiple studies in rodents have shown a connection between the gut microbiota and the central nervous system^8,9^. Microbiota signals and byproducts have been found to play a role in regulating various aspects of brain-related processes and behavior, including stress responses, anxiety and emotional behaviors, cognitive function, myelination, neurogenesis, microglia maturation, and blood-brain barrier integrity^10,11^.

Recent studies also suggest a role of gut microorganisms in neurodevelopmental disorders, including autism spectrum disorders (ASD)^12^. Both patients with ASD and mouse models have been found to have GI dysfunction and changes in the gut microbiota^13–16^. Intriguingly, the transfer of fecal microbiota from human donors with ASD was able to induce hallmark autistic behaviors in the recipient mice, suggesting the existence of a causal link between ASD neurological impairments and the intestinal microbes^17^.

Despite emerging evidence has shown a link between microbiota and neurodevelopmental disorders, no studies have examined the gut-microbiota-brain axis in CDD patients thus far.

Here, for the first time, we analyzed the fecal microbiota composition in a cohort of CDD patients with respect to their healthy neurotypical siblings or relatives. Our data demonstrate the existence of differences in the diversity and composition of the intestinal bacteria in CDD. Moreover, we investigated changes in the microbiota composition in the CDD cohort based on the severity of GI disturbances, frequency of seizures, the presence of sleep disorders, the type of food intake (feeding through food in pieces or smoothies), the impairment of neuro-behavioral features (by the RTT Behaviour Questionnaire), and the possibility of ambulation.

## RESULTS

### Cohort description

We enrolled 17 CDD patients, with identified *CDKL5* gene pathogenic variants (mean age ± SD, 10.9 ± 10.6; 15 females, 2 males). Clinical features are summarized in Table 1, whereas a detailed report on scale scoring is provided in the supplementary Table S1 online. As a control group, we collected stool samples from 17 healthy siblings (or parents if the patient was an only child), who were not on any medication (mean age 28.5 ± 17.2; 13 females).

**Table 1.**
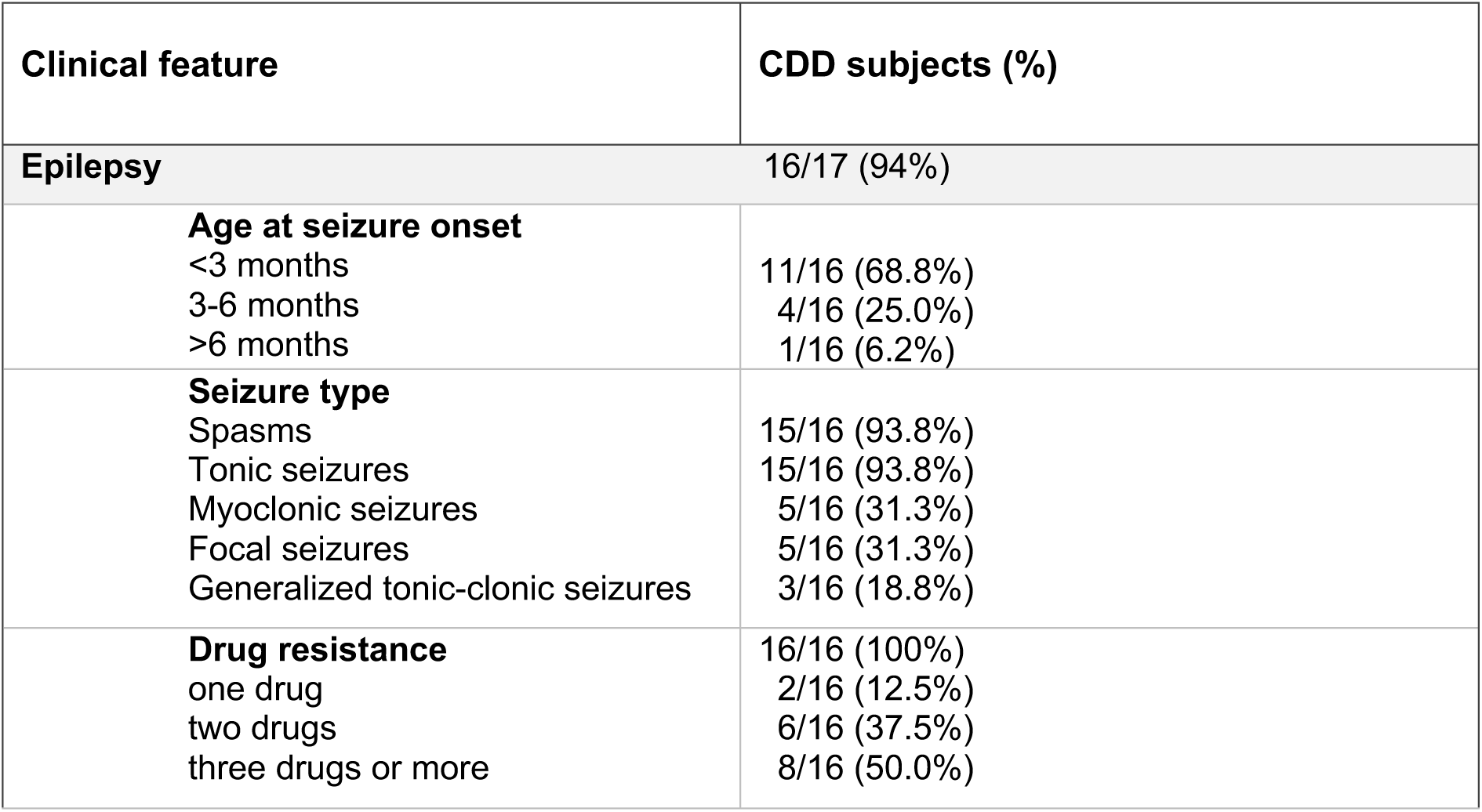
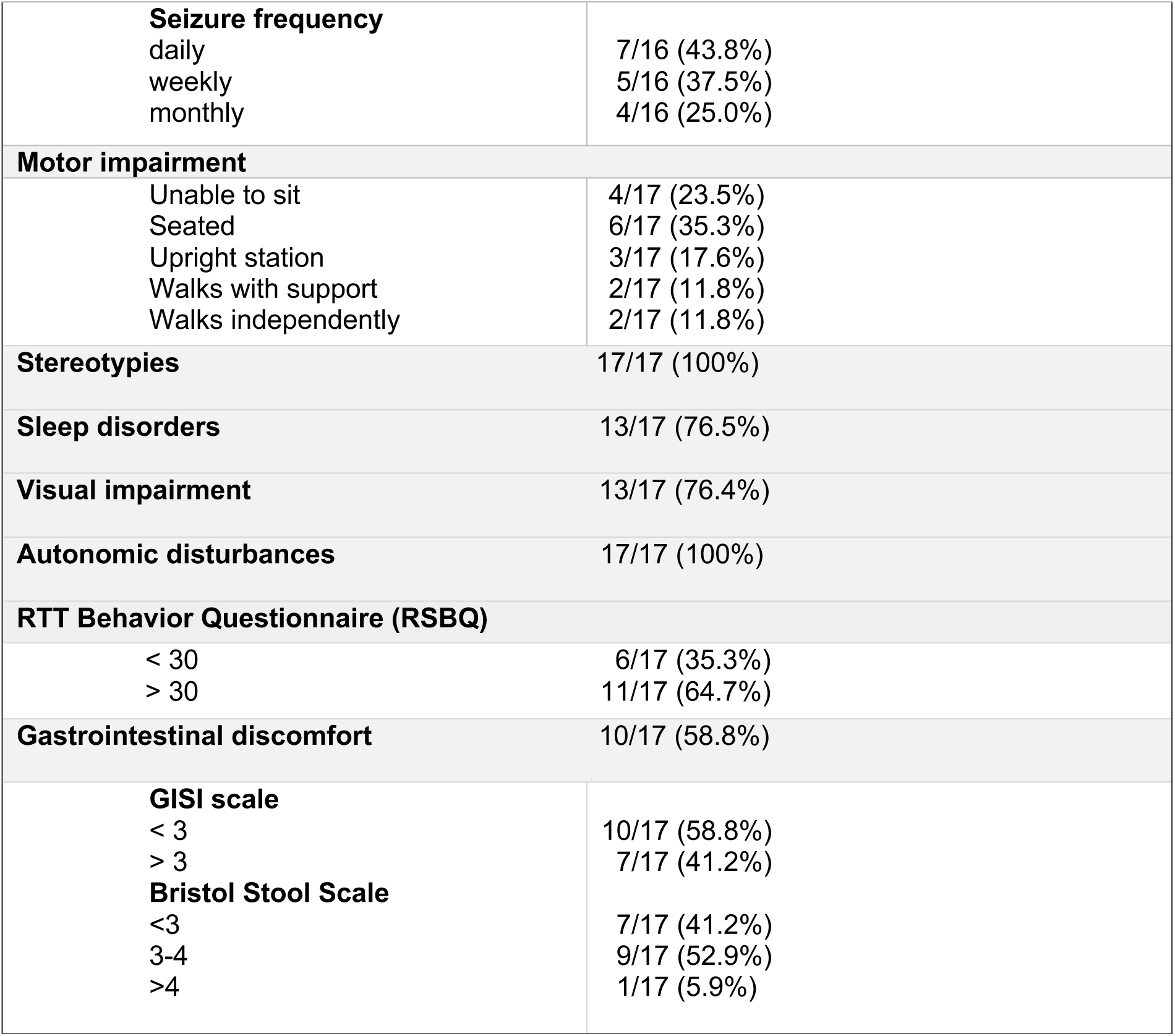
Clinical characteristics of the CDD cohort.

### Dietary nutrient evaluation

Diet is known to be one of the major determinants of the gut microbiota composition, since the types of food we consume directly influence the composition and diversity of microorganisms residing in our GI tract^18^. Despite the 3-days food diary analysis showing a lower daily energy intake in CDD subjects compared to healthy controls (p=0.002), no differences in the percentage of protein, lipid, carbohydrate, and dietary fiber intake in the two groups were observed (Suppl. Fig.1 and Suppl. Table S1). Moreover, the energy intake and macronutrient values are in line with the Italian national recommendations (LARN, Italian Society of Human Nutrition Nutrients and Energy Reference Intake Levels for Italian Population. Available online: https://sinu.it/tabelle-larn-2014/ –last accessed on 27 October 2023-).

**Fig. 1.**
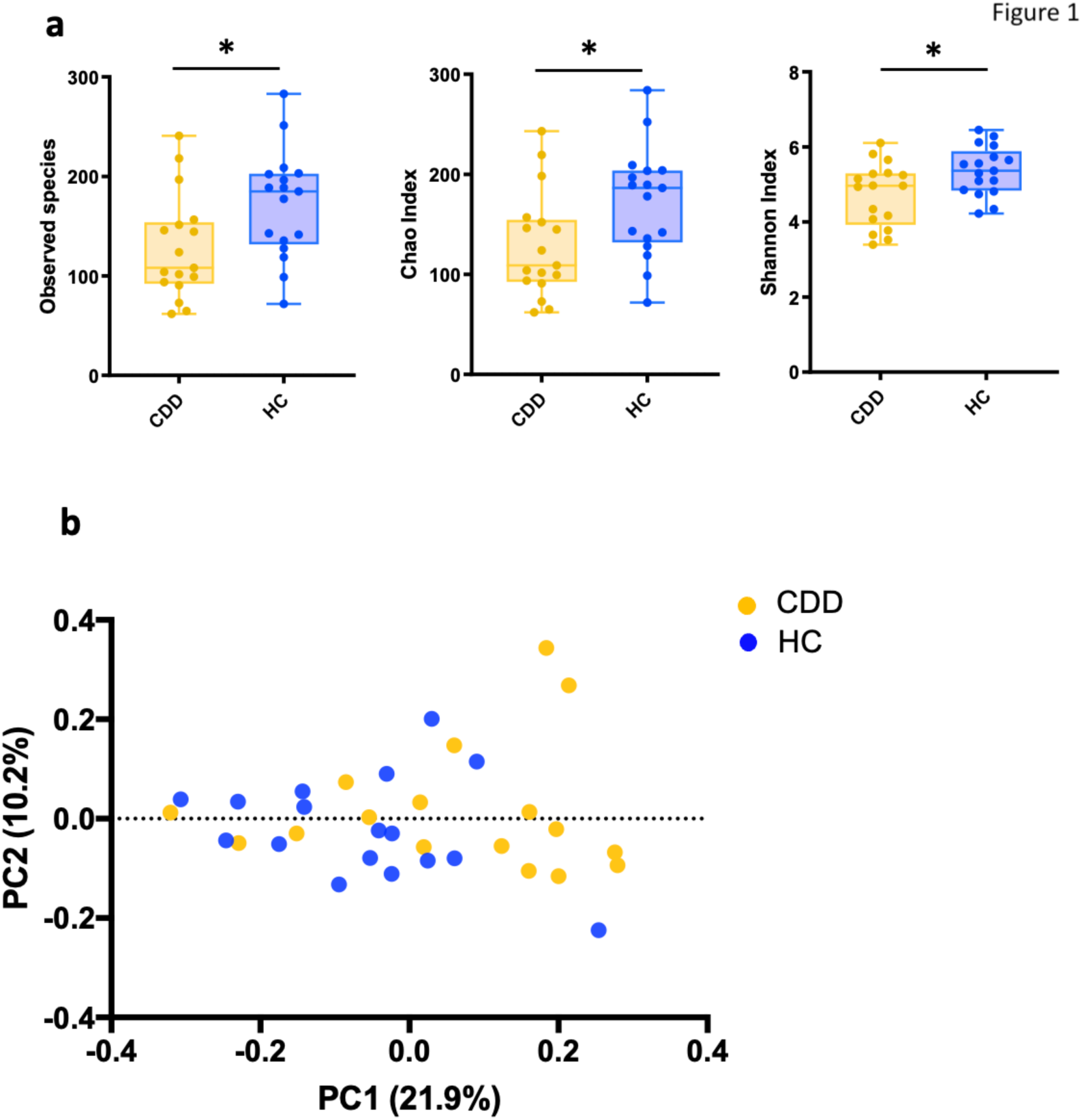
Gut microbial diversity analysis in CDKL5 deficiency patients (CDD) and healthy controls (HC). **a**) Alpha diversity was analyzed through Observed Species, Chao Index and Sahnnon Index metrics. Statistics: Mann-Withney Test, *p value = 0.04. **b)** Principal coordinate analysis of Unweighted Unifrac Distances. Statistics: PERMANOVA test, R = 2.006, p value = 0.014; ANOSIM test, R = 0.098, p value = 0.01. Circles represent single subjects. Error Bars represent SEM.

### Analysis of fecal microbiota composition revealed differences between CDD patients and healthy relatives

The gut microbiota was characterized by next-generation sequencing using V3–V4 hyper-variable 16S rRNA genomic region. On average, 16264,68 ± 4903 high-quality non-chimeric reads were considered for gut microbiota analysis.

We first compared the microbial communities of CDD subjects with those of healthy siblings or parents (Healthy Controls: HC). We found a decrease in biodiversity CDD patients in comparison to healthy controls using three distinct alpha diversity metrics (Figure 1A).

Similarly, the beta-diversity analysis, which assesses the degree of phylogenetic similarity between microbial communities, revealed distinct clustering of the CDD patients’ microbiota compared to the HC group (Figure 1B).

We then explored taxonomic differences in terms of relative abundances in the two groups, CDD and HC. According to the GTDB bacterial taxonomy^19^, at phylum level we observed a significant increase (p=0.006) in Firmicutes (*Bacillales, Erysipelotrichales, Lactobacillales*), and a trend toward a decrease in Firmicutes_A (*Christensenellales, Eubacteriales, Lachnospirales, Oscillospirales*) and in Bacteroidota in CDD patients (Suppl. Dataset 1).

Among the most abundant families (Figure 2A), CDD were enriched in *Erysipelotrichaceae* (p=0.059), *Lactobacillaceae* (p=0.093), *Veillonellaceae* (p=0.1), *Coprobacillaceae* (p=0.107), *Streptococcaceae* (p=0.187), and *Enterococcaceae* (p=0.195). Notably, CDD displayed a significant depletion in CAG-74 (*Christensenellales*) that has been associated with a health status^20^.

**Fig 2.**
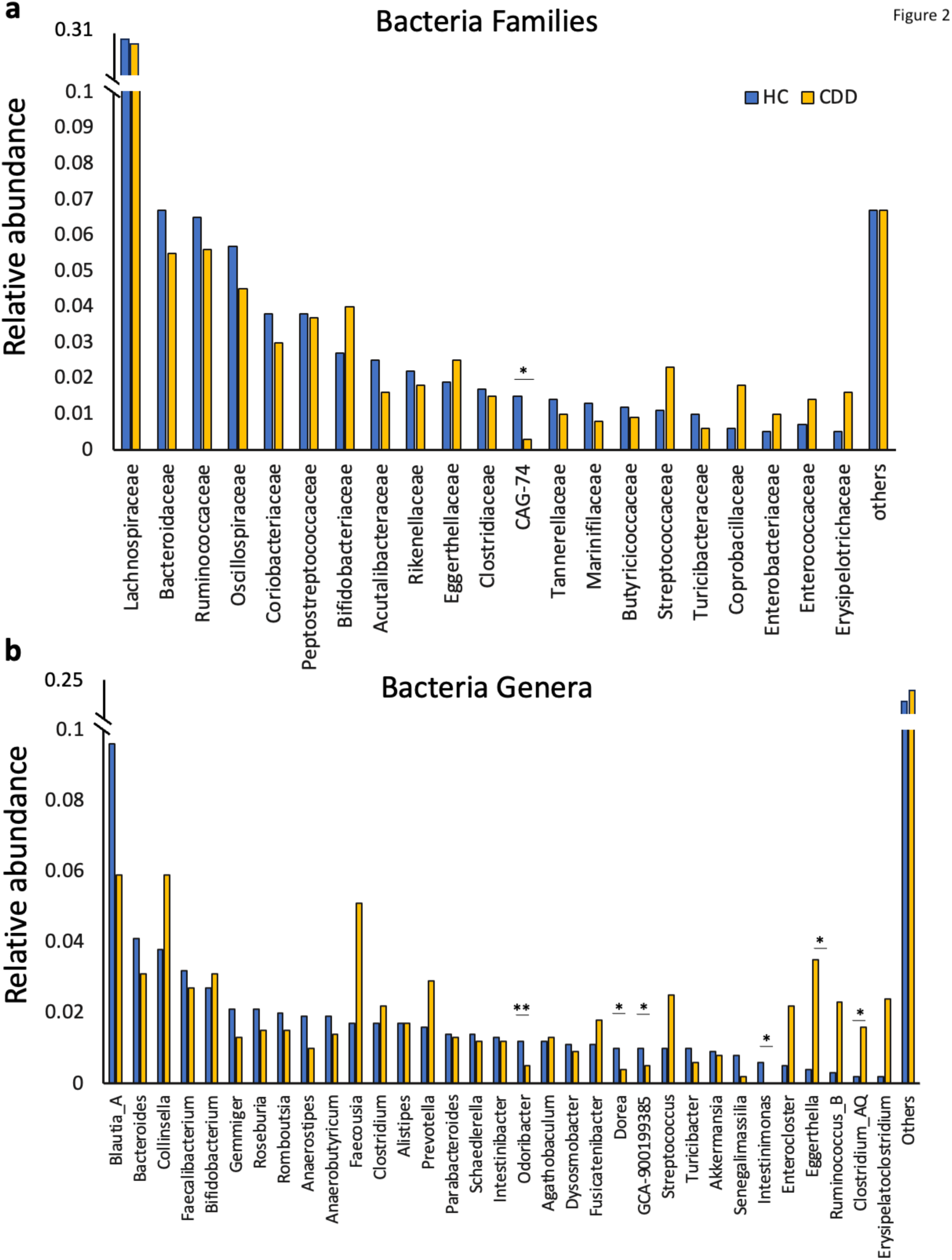
Taxonomy analysis of the gut microbiota in CDD patients and healthy relatives. **a**) Relative abundance of bacteria families, and (**b**) relative abundance of bacteria genera in HC and CDD patients’ fecal microbiota. Statistics: Wilcoxon test, *p value < 0.05, **p value < 0.01. Only families and genera with a relative abundance > .01 in one of the two experimental groups have been reported. See suppl. Dataset 1 for full list and statistics. Others: relative abundance < 0.01.

At genus level (Fig 2B), we found CDD microbial communities to be characterized by an increase in the relative abundance of *Clostridium_AQ* (p= 0.012), *Eggerthella* (p= 0.014), *Streptococcus* (p= 0.114), and *Erysipelatoclostridium* (p= 0.126), and by a decrease in *Eubacterium (Anaerobutyricum*) spp. (p=0.036), *Dorea* (p= 0.004), *Odoribacter* (p= 0.01), *Intestinomonas* (p=0.03), and *Gemmiger* (p= 0.05).

For the complete list of taxa see Suppl. Dataset 1.

Overall, our analysis demonstrates the existence of differences in the composition of the gut microbiota in CDD patients.

### Dissecting the composition of the gut microbiota based on CDD patients clinical features

We then sought to explore differences in the CDD bacterial composition according to each specific clinical feature. In particular, we investigated changes in the microbiota based on the possibility to walk, frequency of seizures, the severity of GI disturbances, the presence of sleep disorders, the score obtained in the RTT Behavior Questionnaire (RSBQ), and the feeding mode (smoothies vs in pieces). Although the number of patients was low, since the disorder is particularly rare, we could observe a tendency to a decrease in alpha-diversity related to the lack of ambulation (Figure 3). Principal coordinate analysis of unweighted unifrac distance showed no significant differences among the CDD patients classified based on clinical features (Suppl. Fig. 2).

**Fig. 3.**
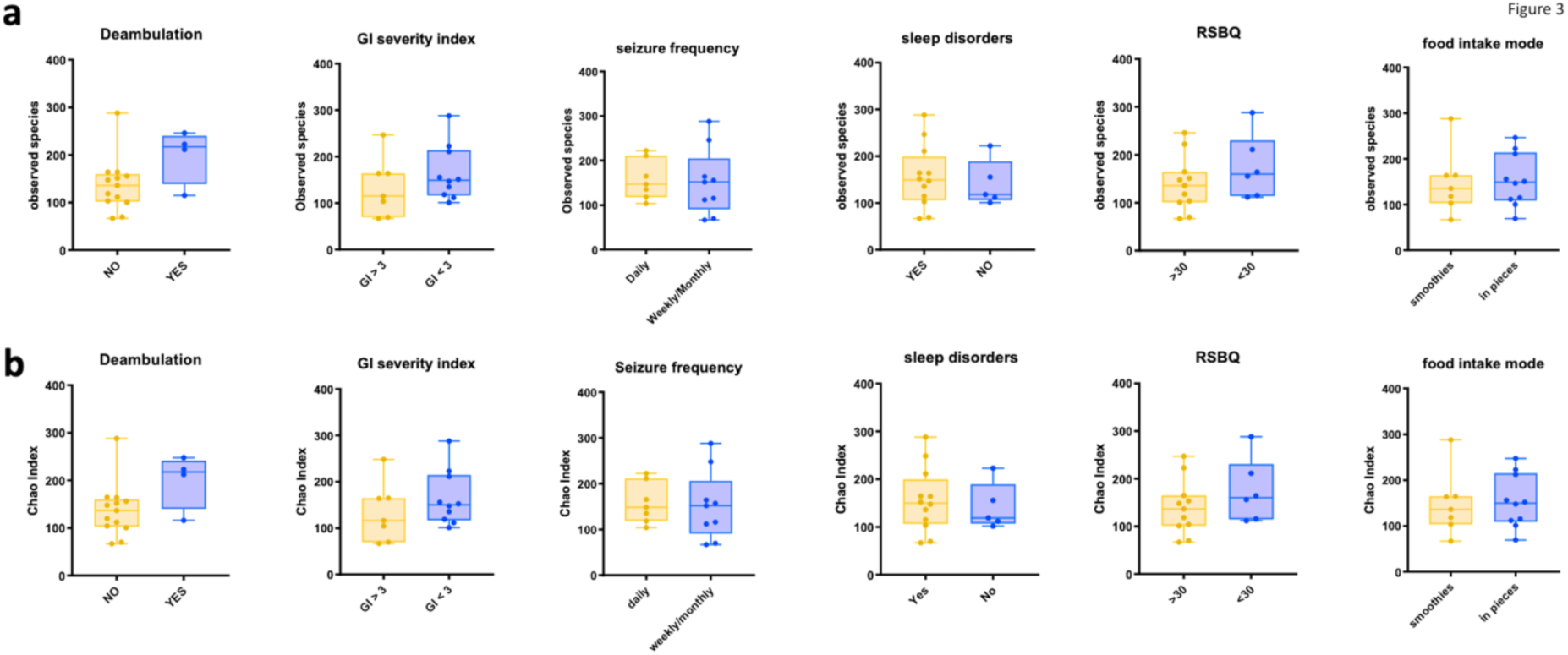
Alpha-diversity in CDD patients grouped according to clinical features. Patients were grouped based on ambulation capacity, GI severity index, the frequency of seizure episodes, the presence of sleep disorders, the score obtained in the RSBQ, and the mode of food intake. **a)** Observed species and **(b)** Chao Index were calculated. Mann Whitney Test showed no significant differences between the tested groups (p>0.05). Circles and squares represent single subjects. Error Bars represent SEM.

The Linear discriminant analysis (LDA) of effect size (LEfSe) method^21^ was used to highlight bacterial taxa that might serve as biomarkers for a defined clinical feature (Figures 4-6).

**Figure 4.**
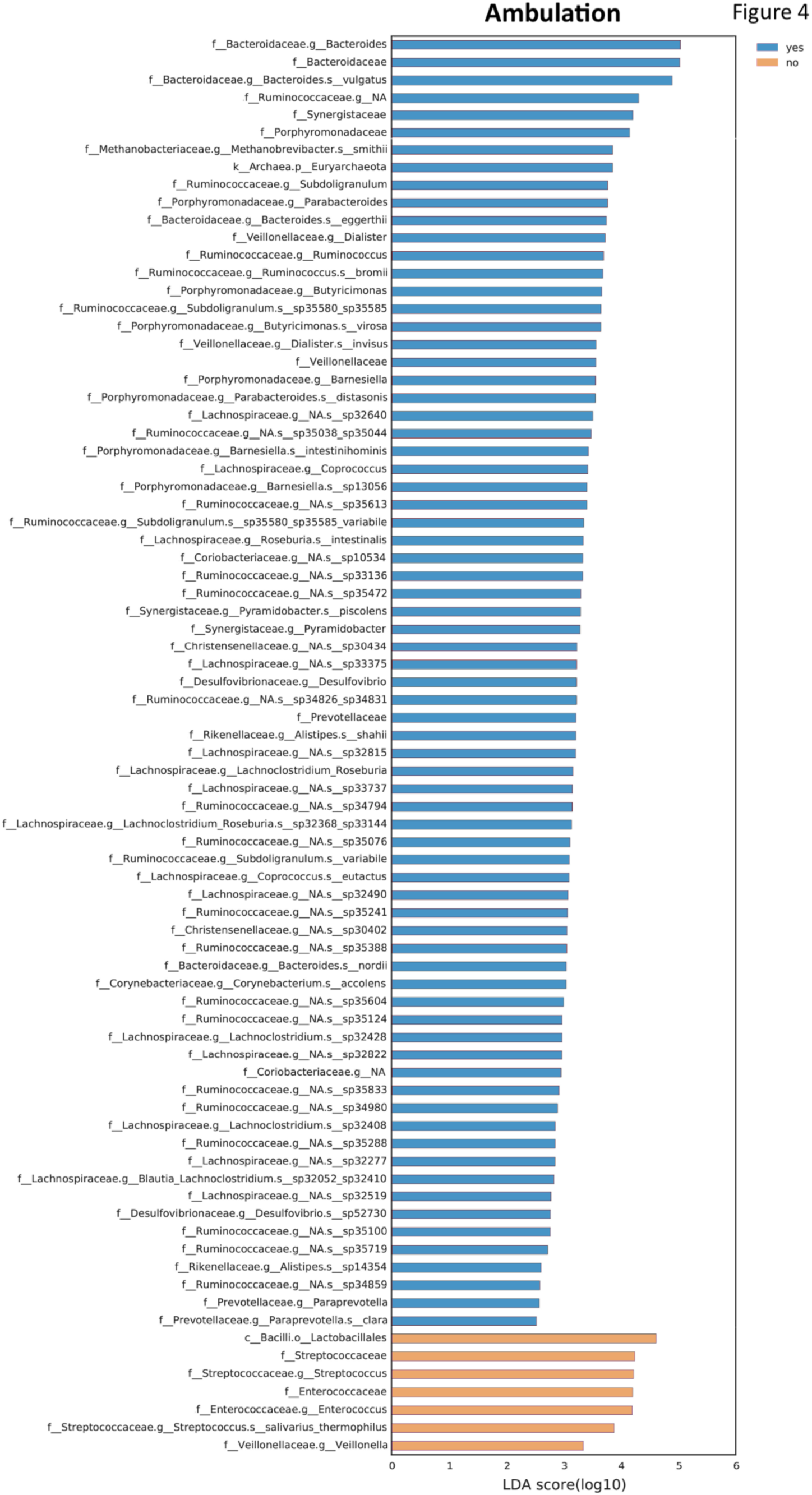
Taxonomic differences in the gut microbiota of CDD subjects according to ambulation. Differently abundant taxa were identified using linear discriminant analysis (LDA) combined with effect size (LEfSe) algorithm. LEfSe analysis identified taxonomic differences in the gut microbiota of CDD subjects according to the capacity of independent walking. Histograms of LDA scores at family level of 16S rRNA gene sequences are shown.

CDD patients not able to deambulate (76.5%) showed a bacterial ecosystem characterized by an increase in the relative abundance of *Enterococcaceae*, *Streptococcaceae* and *Veillonella* spp. (Figure 4). On the contrary, CDD subjects able to walk, either independently or with support, are enriched in *Bacteroidaceae B.vulgatus*), *Methanobacteriaceae* (*M. smithii*), *Ruminococcaceae* (*Subdoligranum*, *Ruminococcus*, and *Roseburia* spp), *Porphyromodanaceae* (*Parabacteroides* spp), and *Dialister* spp.

As refractory epilepsy is a major challenge in CDD, we investigated patients microbiota composition based on seizure frequency. 7 patients were included in the daily seizure group, the other 9 patients showed weekly/monthly epileptic seizures. The LEfSe analysis demonstrated significant differences in the subgroups. Indeed, the daily group had a significant enrichment in bacterial taxa belonging to the family of *Clostridiaceae, Peptostreptococcaceae* (*Romboutsia* spp)*, Erysipelothricaceae,* and *Enterobacteriaceae* (Figure 5b).

**Figure 5.**
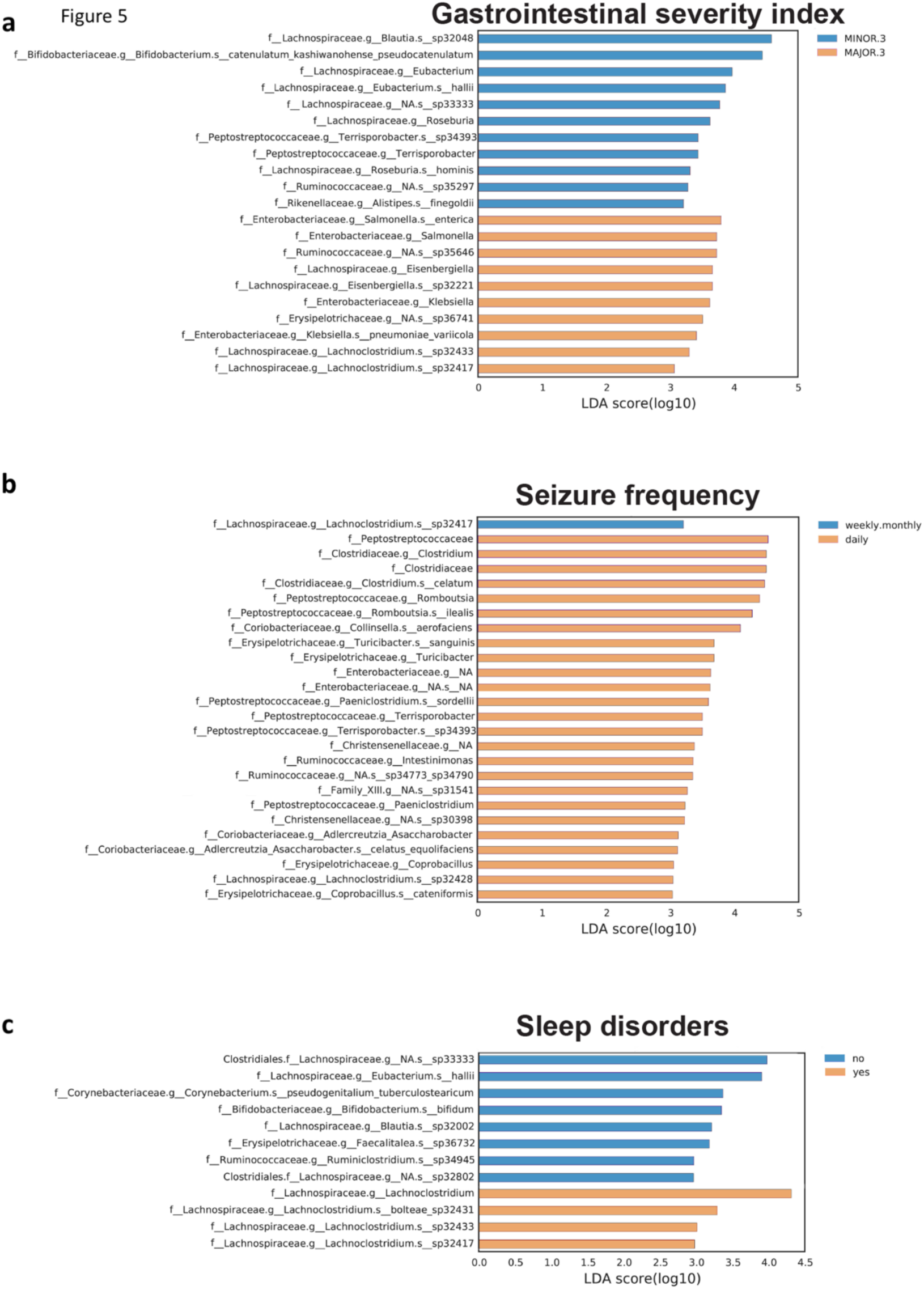
Taxonomic differences in the gut microbiota of CDD subjects according to gastrointestinal symptoms, seizure frequency, and sleep alterations. LEfSe analysis identified taxonomic differences in the gut microbiota of CDD subjects according to the GI severity index (**a**), according to seizure frequency (**b**) and the presence of sleep disturbances (**c**). Histograms of LDA scores at family level of 16S rRNA gene sequences are shown.

Different bacterial taxa were also observed according to GI discomfort evaluated by a modified 6-item GI severity index –GISI-, assessing constipation, diarrhea, stool consistency, stool smell, flatulence, and abdominal pain^22^. Patients displaying a score below 3 (indicating a normal condition or moderate symptoms) showed a significant enrichment in taxa related to GI homeostasis such as *Blautia, Eubacterium, Roseburia*, and *Bifidobacterium* spp. (Figure 5a). On the other hand, subjects scoring above 3 were characterized by an enrichment in *Enterobacteriaceae* that are known to be associated with intestinal inflammation and dysbiosis ^23^.

The genus *Lachnoclostridium* was the indicator taxon for CDD displaying sleep difficulties (Figure 5c). This genus has been associated with intestinal barrier damage and obstructive sleep apnea^24^, was found to be enriched in less physically active older-adult insomnia patients^25^, and negatively correlated with sleep time in a cohort of middle-aged japanese^26^. On the other hand, the gut microbiota in CDD patients not experiencing sleep problems displayed an enrichment of taxa generally related to eubiosis, such as the genera *Eubacterium, Bifidobacterium* and *Blautia* (Figure 5c).

The specific RSBQ scale, evaluating neurological symptoms and intellectual disabilities, was also used for the LEfSe analysis by grouping patients according to their score: >30 or <30 (the higher the score the higher the severity). CDD patients with a score over 30 (64.7%) displayed an enrichment in the genus *Escherichia*, belonging to the *Enterobacteriaceae* family, compared with those with lower scores (<30) (Figure 6a).

**Figure 6.**
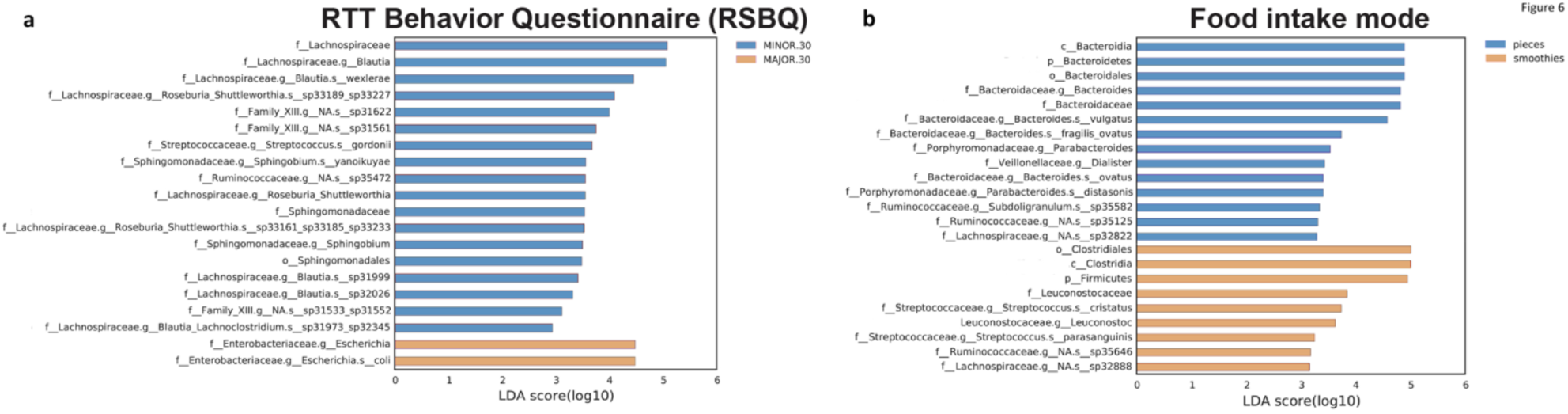
Taxonomic differences in the gut microbiota of CDD subjects related to the RSBQ scale and food intake mode. LEfSe analysis identified taxonomic differences in the gut microbiota of CDD subjects according to the RTT behavior questionnaire (**a**) and according to the food intake mode (**b**). Histograms of LDA scores at family level of 16S rRNA gene sequences are shown.

Significant differences in composition were also observed based on the type of food ingestion type: in pieces *vs* smoothies. We observed an overall enrichment in Bacteroidia linked to food ingestion in pieces (58.8%), and in Firmicutes in subjects eating smoothies (41.2%). *Leuconostocaceae* and *Streptococcaceae* were more present in patients eating smoothies, while the consumption of food in pieces was associated with an enrichment in *Bacteroides, Parabacteroides, Dialister* and *Subdoligranum* spp. (Figure 6b).

Overall, this analysis demonstrates that the grade of clinical features observed in CDD patients is associated with the enrichment specific bacteria taxa, which could potentially serve as candidate biomarkers of symptom severity.

## DISCUSSION

The disruption of the microbiota-gut-brain axis, attributed to disturbances in gut microbiota (i.e., dysbiosis), is considered a potential contributing factor to various neurodevelopmental disorders, including ASD, rare neurological diseases, and epilepsy^8,16,27,28^. Among these disorders, CDD stands out due to its intricate and diverse clinical presentation, encompassing global developmental delay, intellectual disability, speech impairments, visual and motor skill deficits, sleep disturbances, and GI issues^29^. Notably, drug-resistant seizures often manifest early and serve as a primary reason for seeking medical attention^30^, classifying CDD as a developmental and epileptic encephalopathy.

Although GI disturbances are very common among CDD patients and subclinical immune dysregulation has been documented^4^, the relationship between the gut-microbiota-brain axis and CDD has thus far been neglected. In light of these considerations, we conducted a study involving individuals with CDD, as well as their healthy siblings or parents, to investigate the potential connections between alterations in fecal microbiota and the distinct features associated with CDD.

The data obtained in our investigation demonstrate that biodiversity in CDD microbiota is reduced with respect to HC. A decrease in alpha diversity is a phenomenon where the richness of microorganisms within the gut becomes reduced and the community turns uneven. Since richness in bacteria taxa is often associated with a healthy and functional-redundant gut ecosystem, its decrease has been observed in various fields of preclinical and clinical research including inflammatory bowel diseases (IBD), and metabolic disorders. Several studies have explored the relationship between alpha diversity and neuropsychiatric conditions such as ASD^16,31^, however there is still not a clear causal link between alteration in alpha diversity metrics and neurological symptoms. Reduced alpha diversity is a common feature of the gut microbiota in individuals with IBD, which includes conditions like Crohn’s disease and ulcerative colitis. In IBD, the loss of microbial diversity is thought to be linked to chronic inflammation and immune system dysfunction^32^. In light of this evidence, one might hypothesize that the noted reduction in diversity within our CDD patient group could be linked to GI issues and subclinical immune dysfunction. Additionally, the notion that changes in gut microbiota diversity in association with GI problems may contribute to the worsening of neurological symptoms in CDD, potentially impacting the severity of seizures, is conceivable. Nevertheless, it is crucial to conduct further investigations to validate these hypotheses.

Phylogenetic analysis, as revealed by beta diversity assessment, showed a statistical significant clustering of CDD gut microbiota and their relatives, suggesting differences in microbial composition between these two groups. It is worth noting that CDD patients’ intake of macronutrients and dietary fibers was comparable to that of HC. Thus, we can reasonably exclude the possibility that changes in gut microbiota composition are solely linked to the type of diet consumed by CDD patients.

In our previous work on RTT, a disorder sharing several clinical features with CDD, we observed an enrichment in *Bacteroidaceae, Clostridium*, and *Sutterella*, and a reduction in *Prevotella* and *Coprococcus*^5^ in RTT compared to HC. Very similar findings emerged from a study on a larger RTT cohort by Thapa et al., (2021)^33^. The mean relative abundance of *Bacteroides*, *Parabacteroides*, and *Clostridium XlVa* were higher, while *Prevotella* and *Faecalibacterium* were lower in the RTT group compared with the control group.

In the present cohort, compared to healthy siblings, we observed that the main abundant families in CDD were *Erysipelotrichaceae, Lactobacillaceae, Veillonellaceae, Coprobacillaceae, Streptococcaceae, and Enterococcaceae*. Additionally, we found a significant depletion in CAG-74 (*Christensenellales*). Interestingly, the relative abundance of these taxa has been reported as an hallmark for drug-resistant epilepsy^34^, indicating that their presence could be attributed to the specific impact of frequent seizures experienced by CDD subjects.

Collectively, these data suggest a potential link between clinical features of epilepsy in CDD and their fecal microbiota composition.

We also noticed a peculiar increase in the relative abundance of bacterial species characteristic of the oral microbiota in CDD, including *Granulicatella adiacens, Streptococcus salivarius and Veillonella parvula.* Oral-derived microorganisms have been shown to trigger inflammatory responses, promoting their translocation within the intestinal microenvironment^35^, creating a potentially harmful vicious cycle^36^. Intestinal inflammation, as evidenced by recent studies^37^, has been shown to not only promote the occurrence of seizures but also to influence the responsiveness to anti-seizure medication (ASMs)^38^. This may underscore a plausible association between refractory epilepsy in CDD and gut inflammation orchestrated by the microbiota.

Beyond oral-derived bacteria, additional changes in the microbiota suggest the presence of inflammatory triggers originating from microbial sources in CDD. For instance, *Eggerthella* spp, belonging to the phylum Actinobacteria, has been associated with a variety of human inflammatory diseases such as multiple sclerosis and rheumatoid arthritis^39,40^. On the other hand, the CDD gut microbiota is deficient in short chain fatty acids (SCFA)-producing genera such as *Eubacterium*, *Intestinomonas*, *Gemmiger* spp, recently proposed as probiotic species^39,41^. SCFAs, in particular butyrate, exert a protective role enhancing the integrity of the intestinal barrier and dampening gut inflammation^42^.

To better understand the possible contribution of different bacterial taxa in influencing CDD clinical phenotype, we investigated the gut microbiota composition in relation to each symptom severity.

As a developmental epileptic encephalopathy, the most prominent and presenting feature of CDD is the early onset and refractoriness of seizures, with a median onset at 6 weeks of age in 90% of subjects^29,30,43^. In CDD, seizures scarcely respond to conventional ASM, increasing patient and caregiver burden.

It has been suggested that intestinal dysbiosis may contribute to the mechanism of drug-resistant epilepsy^44^. A study published in 2018 reported that people with refractory epilepsy had higher levels of Clostridiales bacteria in their gut microbiome^45^. In our investigation, we found a significant enrichment of the family *Clostridiaceae*, *Peptostreptococcaceae*, *Coriobacteriaceae, Erysipelotrichaceae, Christensenecellaceae* and *Ruminococcaceae*, in CDD patients who experienced daily epileptic seizures. *Collinsella*, belonging to *Coriobacteriaceae,* have been shown to correlate with increased intestinal permeability and to induce the expression of IL-17, a cytokine with a recognized role in epileptogenesis^40,46^. Neuroinflammation is a key player in triggering seizures and in sustaining their occurrence regardless of ASM treatment^47^. A damaged gastrointestinal barrier, a common condition promoted by local inflammation, may lead to bacteria-induced cytokines to directly enter the bloodstream, eventually reaching the brain. Moreover, this scenario can favor the leakage of bacterial products, such as lipopolysaccharides and toxic metabolites, which directly promote systemic inflammation^48^. For instance, clostridia, as spore-forming and toxigenic microorganisms, and *Erysipelotrichaceae,* widely reported as increased in inflammation-related GI disorders^49^, may worsen the chronic inflammatory status within the gut, perpetrating the seizure stimulus.

Severe and extensive motor impairment is another common feature in CDD, primarily because hypotonia is an early characteristic in these children. The achievement of gross motor milestones occurs at a slower pace compared to neurotypical subjects. Independent walking is achieved by only 22–23%, the raking grasp by 49% by the age of 5, and the pincer grasp by a mere 13%^43^.

Regarding the relationship between gut microbiota alterations and motor impairment, previous studies found differences in several genera between children with cerebral palsy and epilepsy (CPE) and healthy controls^50^. Specifically, the CPE group, sharing features with CDD, showed higher proportions of *Bifidobacterium, Streptococcus, Akkermansia, Enterococcus, Prevotella, Veillonella, Rothia*, and *Clostridium IV*. In contrast, the relative abundance of the genera *Bacteroides, Faecalibacterium, Blautia, Ruminococcus, Roseburia, Anaerostipes*, and *Parasutterella* was notably decreased in those patients. In our group of CDD patients, those not walking showed enrichment in the relative abundance of *Enterococcaceae, Streptococcaceae* and *Veillonella*. *Streptococcus* and *Veillonella* often coexist in the gut niche, probably in cross-feeding and metabolic cooperations and have been demonstrated to induce proinflammatory cytokines such as IL-6 and TNF-α, contributing to local inflammation^51^.

GI and nutritional issues are prevalent in CDD patients, with constipation, dysphagia, and GI reflux requiring management in most cases^29^. GI symptoms may vary not only in time and in intensity but also in the clinical features.

In ASD subjects, the analysis of multiple datasets has revealed that numerous taxa discriminate between subjects affected by constipation or affected by diarrhea. These taxa include members of the *Lachnospiraceae* family and species of *Bacteroides, Bifidobacterium, and Streptococcus*^52^. In our previous study in a small Italian RTT cohort, we identified an enrichment of the gut microbiome by two genera, *Bacteroides and Clostridium*^5^. In an American RTT cohort, the overall pattern of the microbiome revealed a higher relative abundance of *Bacteroides spp. and Clostridium spp*. and lower relative abundance of *Prevotella spp. and Faecalibacterium spp.*, suggesting a change in the gut bacterial composition towards an inflammatory community potentially responsible for bowel disease^33^. In this study, the LefSE analysis highlighted an increase in the relative abundance of *Lachnoclostridium* and *Enterobacteriaceae* in CDD patients with the most severe GI symptoms. This finding suggests, once again, that inflammation serves as a mediator of the clinical features within the microbiota and gut homeostasis.

Regarding the management of feeding difficulties, CDD patients are forced to change their dietary habits, and strategies for facilitating food intake are frequently adopted by caregivers. Indeed, in the enrolled CDD group, 7 out of 17 subjects consumed a smoothie diet. Studies on the gut microbiota have indicated that diet plays a significant role in shaping the composition of intestinal microbes^53^. Changes in dietary fiber intake, food processing methods, and food consistency can influence the types and proportions of microorganisms residing in the GI tract. Despite no difference in the diet macronutrient intake, we observed that CDD patients consuming a blended diet had an increase in *Firmicutes*, whereas those ingesting food in pieces were characterized by a higher relative abundance in *Bacteroidia*. These findings are in agreement with a previous study comparing gastrostomy-fed children with blended “homemade” diets and their siblings^54^.

Sleep difficulties have often been reported in CDD patients, and appear to be more common in CDD than in RTT^1,55–57^. Importantly, sleep disturbances represent a considerable source of concern for caregivers and affect the overall quality of life for CDD patients and their families.

In our cohort, most CDD subjects experienced a sleep problem (76%), in line with the data reported by Mangatt and coworkers describing that, during their lifetime, CDD patients may show current night waking, sleep breathing disorders, disorders of arousal, sleep-wake transition, and excessive diurnal somnolence^55^.

To date, only a few studies have explored the gut microbiota alterations in sleep disorder comorbidity in neurodevelopmental disorders. Specifically, investigations in patients with ASD and sleep disorders have found that *Faecalibacterium* and *Agathobacter* abundance is reduced with respect to ASD children without sleep disorders^58^. Both these genera produce butyrate, which is a SCFA that plays an important role in gut physiology^59^. The decrease in butyrate and its metabolites may determine an increase in serotonin and a decrease in melatonin, potentially aggravating sleep problems^60^. Moreover, in a recent study exploring insomnia in patients with major depression, *Coprococcus* and *Intestinibacter* were associated with sleep quality, independent of the severity of depression. *Coprococcus* is a butyrate-producer itself, while *Intestinibacter* may act via immune inflammatory mechanisms or carbohydrate metabolism^61^. In our analysis, the fecal microbiota in CDD patients experiencing sleep disturbances was characterized by a significant enrichment in the family *Lachnospiraceae*, genus Lachnoclostridium. On the other hand, the microbiota in patients with a normal sleep pattern was enriched in the genera *Eubacterium*, *Bifidobacterium* and *Blautia* which are associated with eubiosis. *Bifidobacterium* shows SCFA-producing ability, as well as *Eubacterium hallii* which is a producer of butyrate and propionate, and is able of metabolizing glycerol to 3-hydroxypropionaldehyde (3-HPA), with antimicrobial properties^62^. Based on our analysis, there appears to be an association between sleep disturbances in CDD patients and a potential decrease in SCFA-producing bacteria. This observation could pave the way for new intervention strategies, such as targeting the microbiota with probiotic bacteria enriched in SCFA-producing species, to improve both GI abnormalities and sleep quality.

Finally, to evaluate the contribution of microbial alterations to neurological and behavioral symptoms, we stratified patients based on RSBQ score, a caregiver-completed scale. This questionnaire is among the most extensively utilized neuro-behavioral assessment tools in clinical studies related to RTT, primarily due to its specific focus on the core features of the syndrome^63^. We categorized CDD patients as having higher clinical severity when their RSBQ score was >30 points^64^.

In our cohort, CDD patients with a score over 30 displayed an enrichment in the genus *Escherichia*, belonging to the *Enterobacteriaceae* family, compared with those with lower scores (<30). Multiple studies demonstrate the association of this family with dysbiosis in pathological conditions such as IBS^65^. As a consequence of or a potential promoting factor of GI disturbances, we might speculate that the presence of a low grade-inflammatory microenvironment in CDD gut could favor the blooming of *Enterobacteriaceae,* further unbalancing intestinal homeostasis^66^. As the gut and the brain are continuously communicating through the gut-brain axis, an enrichment of potentially dysbiotic bacteria (i.e. *Enterobacteriaceae*) might interfere with this crosstalk system, leading to worsen neurological outcomes and behavior. Indeed, speculative conclusions about microbiota-dependent modifications in brain function and behavior have been raised, specifically in relation to subjects with ASD. In humans, new data suggests the link between behavioral/neurological symptoms and the gut microbiota in ASD, demonstrating that an alteration of Bacteroides with multiple *Bifidobacterium longum* subsp longum, swine fecal bacteria SD Pec10, *Ruminococcus* sp N15MGS 57, and *Parabacteroides merdae* could be associated to social and emotional/behavioral symptoms^67^.

In line with the above mentioned data, a plethora of investigations in mouse models have demonstrated the involvement of the gut microbiota in behavioral outcome and brain plasticity^68–70^. Gut bacteria could modulate stress responses^71^, anxiety-like behavior^72^, social behavior^73–75^, fear conditioning^76^, learning and memory^77,78^. Intriguingly, in ASD rodent models the gut microbiota and some of its metabolites could modulate autistic-like behavior^13,15,17^. Mechanisms related to specific probiotic action on vagus nerve, hypothalamic oxytocin signaling pathway or the increase in the level of bacteria-derived metabolites have been proposed^13–15^. While these findings in preclinical models offer intriguing insights, it is essential to recognize their preliminary nature, indicating the need for additional research to ascertain their translational relevance in humans.

## Conclusion

The microbiota-gut-brain axis has been suggested to play a role in several neurodevelopmental disorders and recent data support its involvement in drug-refractory epilepsy. For the first time, our study sheds light on the potential connection between gut microbiota alterations and the multifaceted CDD phenotype. Microbiota alterations may contribute to the most common disease comorbidities, mainly by promoting a subclinical inflammation state. Indeed, most of the increased/reduced bacterial taxa have a demonstrated role in inducing/attenuating gut inflammation. Further research is needed to explore the mechanisms underlying these connections and to develop targeted interventions for improving the wellbeing of individuals with CDD. Finally, as suggested by the specific changes observed in certain bacteria families and genera, intestinal microbes may become a novel biomarker in CDD. However, it is important to note that while these findings are promising, further research is necessary to establish the microbiota’s validity as a biomarker for CDD. Longitudinal studies involving larger patient cohorts and more extensive clinical data will be crucial in confirming these associations and exploring the potential of microbiota-based diagnostics and monitoring tools. Ultimately, this approach could have a transformative impact on the diagnosis and management of CDD, offering new insights into its pathophysiology and potential therapeutic interventions.

## METHODS

### Cohort recruitment

In collaboration with the Italian Association of CDKL5 families (“Insieme verso la cura”), we recruited individuals who had been diagnosed with CDKL5 disorder and had a confirmed pathogenic variant in the *CDKL5* gene. As a control group, we included healthy siblings or parents (for only child) of the affected individuals, who were not currently taking any medications. We excluded individuals who had used antibiotics or probiotics within the three months prior to the study.

At the enrollment, caregivers were asked to fill a 3-day dietary survey. The diary included three consecutive days, one of which was during the weekend. Dietary food records were processed using a commercially available software (MètaDieta, METEDA srl, Italy). Anthropometric evaluation completed the nutritional survey.

CDD patients underwent a thorough characterization process, which involved comprehensive clinical evaluations. This evaluation encompassed a detailed examination of their medical history and physical condition, including an assessment of neurological and behavioral features, as well as the identification of any comorbidity. The overall clinical assessment was measured by the Rett Syndrome Behavioral Questionnaire (RSBQ)^79^, considered specific for CDD and RTT syndromes. Additionally, a GI assessment was conducted to gather information about the individual’s GI health. GI discomfort was evaluated by the Gastrointestinal Severity Index (GISI)^22^, and stool transition time was estimated by the Bristol Stool Form Scale (BSFS, Lewis and Heaton, 1997).

Epilepsy data were also recorded, in particular: age at seizure onset, seizure characteristics at onset and at follow-up, anti-seizure medication history and current treatment. Seizure frequency at the time of the study was recorded using seizure diaries.

### Fecal DNA extraction and 16S rRNA sequencing

Bacterial DNA was extracted from fecal samples using the QIAamp Powerfecal DNA kit (Qiagen, Germany), and its concentration was quantified by Qubit® (Thermo Fisher Scientific, Waltham, WA).

The 16S rRNA sequencing and analysis was performed in service by Zymo Research (Germany). Briefly, DNA samples were prepared for targeted sequencing with the Quick-16S™ NGS Library Prep Kit (Zymo Research) and the Quick-16S™ Primer Set V3-V4 (Zymo Research). The sequencing library was prepared using an innovative library preparation process in which PCR reactions were performed in real-time PCR machines to control cycles and therefore limit PCR chimera formation. The final PCR products were quantified with qPCR fluorescence readings and pooled together based on equal molarity. The final pooled library was cleaned up with the Select-a-Size DNA Clean & Concentrator™, then quantified with TapeStation® (Agilent Technologies, Santa Clara, CA) and Qubit® (Thermo Fisher Scientific, Waltham, WA).

The ZymoBIOMICS® Microbial Community DNA Standard (Zymo Research, Irvine, CA) was used as a positive control for each targeted library preparation. Negative controls (i.e., blank library preparation control) were included to assess the level of bioburden carried by the wet-lab process.

The final library was sequenced on Illumina® MiSeq™ with a v3 reagent kit (600 cycles). The sequencing was performed with 10% PhiX spike-in.

### Bioinformatics Analysis

Amplicon sequence variants (ASVs) were identified from 16S paired-end sequencing using the Divisive Amplicon Denoising Algorithm (DADA2) pipeline, including filtering and trimming of the reads (version 1.16.0)^80^. Reads per sample were trimmed to 10900 reads in order to compensate for the sequencing unevenness of the samples and to provide a consistent minimum amount for the downstream analysis, carried out through the “phyloseq” package (version 1.34.0)^81^. Alpha-diversity evaluation was performed according to several microbial diversity metrics (i.e., chao1, Shannon Index, observed species). Beta-diversity analysis was conducted using unweighted Unifrac metrics^82^ and through the principal coordinates analysis (PCoA).

Taxonomy was assigned to the ASVs using the 8-mer-based classifier from the 11.5 release of the RDP database^83^ and using the GTDB 16S rRNA database (release r207)^84^. In the analysis of the gut microbiota in CDD patients based on clinical features, taxonomies that have significant abundance among different groups were identified by LEfSe^21^ using default settings.

#### Statistical Analysis

Comparisons were performed using Prism software Prism version 7 (GraphPad Software, San Diego, CA, USA) or Python.

Alpha-diversity between CDD and HC groups, or intra CDD groups based, was tested through Mann–Whitney U-test.

Beta-diversity: to test whether two groups of samples were significantly different, analysis of similarities (ANOSIM) or permutational multivariate ANOVA (PERMANOVA) were calculated using the Python library scikit-bio (http://scikit-bio.org/).

For evaluating differences in relative abundances of bacterial groups (Families and Genera) between CDD and HC, a Wilcoxon test was performed.

Due to the small number of samples (17 CDD vs 17 HC), no correction methods have been applied. N’s represent single subjects unless otherwise stated. Statistical significance was defined in the figure panels as follows: \**P* < 0.05, \*\**P* < 0.01.

## Acknowledgements

The authors are extremely thankful to the families and the children of the association “Insieme verso la cura’’ for their participation and support in the study.

We thank to Prof. Tommaso Pizzorusso (Scuola Normale Superiore, Pisa) for his critical insights on the manuscript and figures. We thank Dr. Raffaele Mazziotti (University of Florence, Italy) for his help with statistics.

Special thanks to Andrea Tognozzi and to all the members of Tognini’s Lab for comments on figure style.

## Author contributions statement

EB, PT and AV designed the study. IV and AV performed subject enrollment and AV analyzed clinical data. OX and EO carried out the sample collection and experiments. EO performed diet analysis. CC, EB, and PT analyzed, and interpreted the data. EB and PT prepared the figures. EB, PT and AV wrote the manuscript. All authors discussed the results and commented on the manuscript. All authors contributed to the article and approved the submitted version.

## Data Availability

16S rRNA-seq data will be available in a public repository upon manuscript acceptance.

## Ethics approval

The study has been approved by the Ethic Committee of ASST Santi Paolo Carlo Hospital, Milano, Italy (2021/EM/317).

