## Supplementary tables and Figures for "Gut microbiota profile in CDKL5 deficiency disorder patients as a potential marker of clinical severity"

**Table S1-** Nutritional values of the enrolled subjects

| | CDD (mean ± SD) | | HD (mean ± SD) | | *p*-Value (CDD *vs* HC) | | Reference Values (LARN) | |
|---|---|---|---|---|---|---|---|---|
| **Variable** | **Children** | **Adults** | **Children** | **Adults** | **Children** | **Adults** | **Children** | **Adults** |
| Energy (kcal) | 1065.4 ± 682.4 | 1280.7 ± 107.0 | 1504.6 ± 405.5 | 1630.9 ± 275.3 | 0.02 | 0.10 | 750-1860 kcal/day | 1240-1910 kcal/day |
| Proteins (g) | 41.9 ± 14.3 | 54.8 ± 4.8 | 51.8 ± 18.8 | 74.8 ± 16.2 | 0.35 | 0.10 | 9-50 (g/day)* | 43-50 (g/day)* |
| Proteins (%E) | 16.5 ± 3.9 | 17.6 ± 0.4 | 13.7 ± 1.4 | 19.0 ± 3.4 | 0.12 | 0.48 | 0.72-0.82 (g/kg/day)* | 0.71 %* |
| Total Carbohydrates (g) | 155.1 ± 1 39.1 | 186.3 ± 21.7 | 205.8 ± 35.2 | 177.0 ± 53.4 | 0.01 | 0.73 | | |
| Total Carbohydrates (%E) | 49.8 ± 12.0 | 56.0 ± 5.2 | 53.54 ± 6.8 | 41.8 ± 9.6 | 0.35 | 0.10 | 45%-60% E** | 45%-60% E** |
| Sugars (g) | 40.0 ± 17.5 | 50.4 ± 7.9 | 76.0 ± 21.0 | 54.1 ± 33.1 | <0.01 | 0.86 | <15% E | |
| Starch (g) | 51.2 ± 82.1 | 47.0 ± 25.8 | 48.5 ± 16.0 | 60.0 ± 42.0 | 0.07 | 0.73 | | |
| Fiber (g) | 12.7 ± 13.7 | 17.6 ± 5.0 | 13.0 ± 6.6 | 15.8 ± 8.6 | 0.21 | 0.86 | 8.4 g/1000 Kcal | 12.6-16.7 g/1000 Kcal |
| Soluble Fiber (g) | 1.2 ± 0.8 | 1.9 ± 0.6 | 1.5 ± 1.0 | 1.2 ± 0.9 | 0.44 | 0.21 | | |
| Insoluble Fiber (g) | 2.9 ± 2.1 | 5.2 ± 2.3 | 4.2 ± 3.7 | 4.8 ± 4.5 | 0.49 | 0.73 | | |
| Fats (g) | 37.3 ± 12.1 | 37.6 ± 6.1 | 43.2 ± 5.7 | 58.9 ± 15.5 | 0.31 | 0.04 | 20%-40% E** | 20%-40% E** |
| Saturated Fats (g) | 12.8 ± 6.2 | 10.5 ± 3.6 | 17.7 ± 4.0 | 22.5 ± 3.6 | 0.20 | <0.01 | <10% E | <10% E |

CDD, CDKL5-deficiency disorder; HC, heathy control; E, energy; *AR, Average Requirement; **RI, recommended intake.

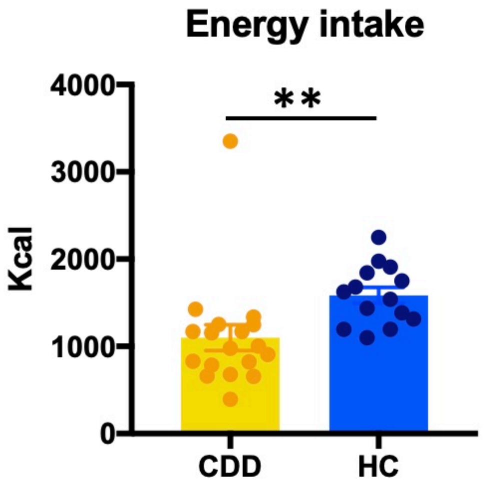


**Suppl. Fig. 1. Food diary analysis.** The enrgy intake quantified as Kcal consumed/day was significantly decreased in CDD patients with respect to HC (Unpaired t-test, **p=0.002). The percentage of lipid, protein, carbohydrate and fiber consumed/day were not different. Circles represent single subject. Error bars represent SEM

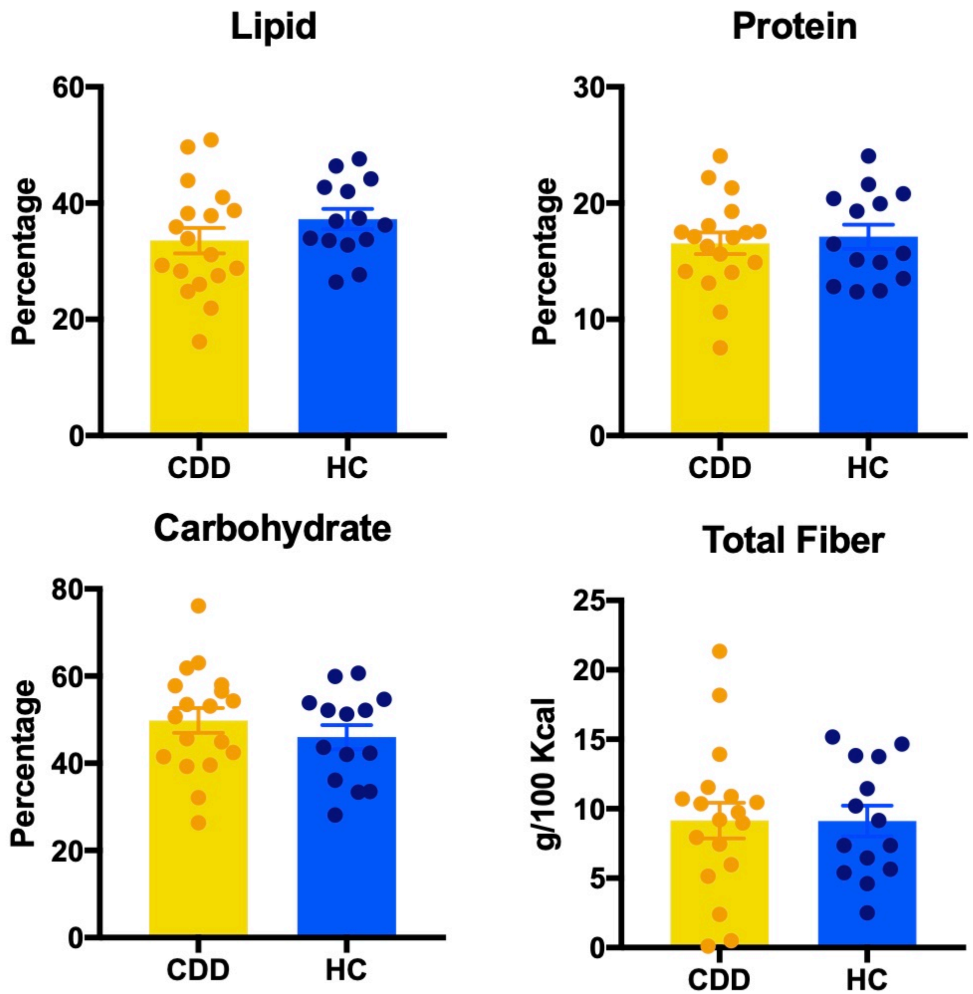

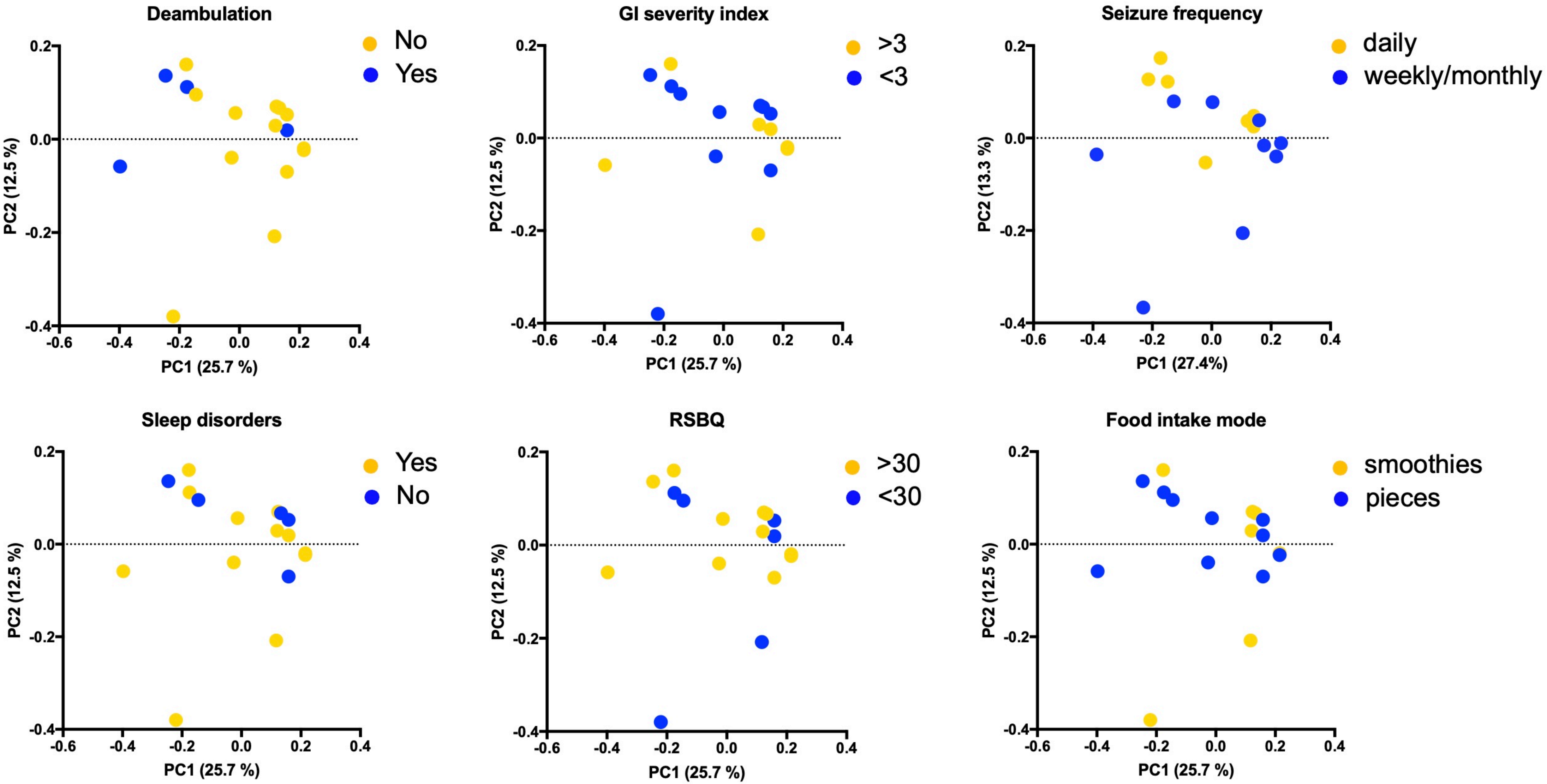


**Suppl. Fig. 2. Beta-diversity in CDD patients.** Patients were grouped based on deambulation capacity, GI severity index, the frequency of seizure episodes, the presence of sleep disorders, the score obtained in the RSBQ, and the mode of food intake. Principal coordinate analysis of Unweighted Unifrac Distances.
Circles represent single subjects.
